# Multi-SKAT: General framework to test for rare variant association with multiple phenotypes

**DOI:** 10.1101/229583

**Authors:** Diptavo Dutta, Laura Scott, Michael Boehnke, Seunggeun Lee

## Abstract

In genetic association analysis, a joint test of multiple distinct phenotypes can increase power to identify sets of trait-associated variants within genes or regions of interest. Existing multi-phenotype tests for rare variants make specific assumptions about the patterns of association with underlying causal variants and the violation of these assumptions can reduce power to detect association. Here we develop a general framework for testing pleiotropic effects of rare variants on multiple continuous phenotypes using multivariate kernel regression (Multi-SKAT). Multi-SKAT models effect sizes of variants on the phenotypes through a kernel matrix and performs a variance component test of association. We show that many existing tests are equivalent to specific choices of kernel matrices with the Multi-SKAT framework. To increase power of detecting association across tests with different kernel matrices, we developed a fast and accurate approximation of the significance of the minimum observed p-value across tests. To account for related individuals, our framework uses random effects for the kinship matrix. Using simulated data and amino acid and exome-array data from the METSIM study, we show that Multi-SKAT can improve power over single-phenotype SKAT-O test and existing multiple phenotype tests, while maintaining type I error rate.

**Grant Numbers:** This research was conducted with support from the following grants from National Institute of Health:

- R01 HG008773 (D.D. and S.L.)
- R01 LM012535 (D.D. and S.L.)
- R01 HG000376 (L.S. and M.B.)
- U01 DK062370 (L.S. and M.B.)

## Introduction

Since the advent of array genotyping technologies, genome-wide association studies (GWAS) have identified numerous genetic variants associated with complex traits. Despite these many discoveries, GWAS loci explain only a modest proportion of heritability for most traits. This may be due, in part, to the fact that these association studies are underpowered to identify associations with rare variants(Korte and Farlow, 2013). To identify such rare variant associations, gene- or region-based multiple variant tests have been developed(Lee et al., 2014). By jointly testing rare variants in a target gene or region, these methods can increase power over a single variant test and are now used as a standard approach in rare variant analysis.

Recent GWAS results have shown that many GWAS loci are associated with multiple traits (Solovieff et al., 2013). Nearly 17% of variants in National Heart Lung and Blood Institute (NHLBI) GWAS categories are associated with multiple traits (Sivakumaran et al., 2011). For example, 44% of autoimmune risk single nucleotide polymorphisms (SNPs) have been estimated to be associated with two or more autoimmune diseases (Cotsapas et al., 2011). Detecting such pleiotropic effects is important to understand the underlying biological structure of complex traits. In addition, by leveraging cross-phenotype associations, the power to detect trait-associated variants can be increased.

Identifying the cross-phenotype effects requires a suitable joint or multivariate analysis framework that can leverage the dependence of the phenotypes. Various methods have been proposed for multiple phenotype analysis in GWAS (Ferreira and Purcell, 2009; Huang et al., 2011; Zhou and Stephens, 2014; Ried et al., 2012; Ray et al., 2016). Extending them, several groups have developed multiple phenotype tests for rare variants (Wang et al., 2015; Broadaway et al., 2016; Wu and Pankow, 2016; Lee et al., 2016; Sun et al., 2016; Maity et al., 2012; Yan et al., 2015; Zhan et al., 2017). For example, Wang et al. (2015) proposed a multivariate functional linear model (MFLM); Broadaway et al. (2016) used a dual-kernel based distance-covariance approach to test for cross phenotype effects of rare variants by comparing similarity in multivariate phenotypes to similarity in genetic variants (GAMuT)(Chiu et al., 2017); Wu et al. (Wu and Pankow, 2016) developed a score based sequence kernel association test for multiple traits, MSKAT, which has been shown to be similar in performance to GAMuT(Broadaway et al., 2016); and Zhan et al. (2017) proposed DKAT, which uses the dual kernel approach as in GAMuT but provides more robust performance when the dimension of phenotypes is high compared to the sample size.

Despite these developments, existing methods have important limitations. Most methods were developed under specific assumptions regarding the effects of the variants on multiple phenotypes, and hence lose power if the assumptions are violated (Ray et al., 2016). For example, if genetic effects are heterogeneous across multiple phenotypes, methods assuming homogeneous genetic effects can lose a substantial amount of power. Although there has been a recent attempt to combine analysis results from different models (Zhan et al., 2017), no scalable methods have been developed to evaluate the significance of the combined results in genome-wide scale analysis. In addition, most existing methods and software cannot adjust for relatedness between individuals; thus, to apply these methods, related individuals must be removed from the analysis to maintain type I error rate. For example, in the METabolic Syndrome In Men (METSIM) study ~ 15% of individuals are estimated to be related up to the second degree.

Here, we develop Multi-SKAT, a general framework that extends the mixed effect model-based kernel association tests to a multivariate regression framework while accounting for family relatedness. Mixed effect models have been widely used for rare-variant association tests. Popular rare variant tests such as SKAT(Wu et al., 2011) and SKAT-O(Lee et al., 2012b) are based on mixed effect models. By using kernels to relate genetic variants to multiple continuous phenotypes, Multi-SKAT allows for flexible modeling of the genetic effects on the phenotypes. The idea of using kernels for genotypes and phenotypes were previously used by the dual kernel approaches such as GAMuT and DKAT. However, in contrast to these two similarity-based methods, Multi-SKAT is multivariate regression based and hence provides a natural way to adjust for covariates and also can account for sample relatedness by incorporating random effects for the kinship matrix. Many of the existing methods for multiple phenotype rare variant tests can be viewed as special cases of Multi-SKAT with particular choices of kernels. Furthermore, to avoid loss of power due to model misspecification, we develop computationally efficient omnibus tests, which allow for aggregation of tests over several kernels and provide fast p-value calculation (Demarta and McNeil, 2005).

The article is organized as follows: in the first section, we present the multivariate mixed effect model and kernel matrices. We particularly focus on the phenotype-kernel and describe omnibus procedures that can aggregate results across different choices of kernels and kinship adjustment. In the next section we describe the simulation experiments that clearly demonstrate that Multi-SKAT tests have increased power to detect associations compared to existing methods like GAMuT, MSKAT and others in most of the scenarios. Further we applied Multi-SKAT to detect the cross-phenotype effects of rare nonsynonymous and protein-truncating variants on a set of nine amino acids measured on 8,545 Finnish men from the METSIM study.

## Material and Methods

### Single-phenotype region-based tests

To describe the Multi-SKAT tests, we first present the existing model of the single-phenotype gene or region-based tests. Let *y_k_* = (*y*_1*k*_, *y*_2*k*_, ⋯, *y_nk_*)*^T^* be an *n* × 1 vector of the *k^th^* phenotype over *n* individuals; X an *n* × *q* matrix of the *q* non-genetic covariates including the intercept; *G_j_* = (*G*_1*j*_, ⋯, *G_nj_*)*^T^* is an *n* × 1 vector of the minor allele counts (0, 1, or 2) for a binary genetic variant *j*; and *G* = [*G*_1_, ⋯, *G_m_*] is an *n* × *m* genotype matrix for *m* genetic variants in a target region. The regression model shown in equation (1) can relate *m* genetic variants to phenotype *k*,

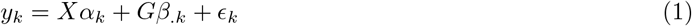

where *α_k_* is a *q* × 1 vector of regression coefficients of *q* non-genetic covariates, *β_.k_* = (*β*_1*k*_, ⋯, *β_mk_*)*^T^* is an *m* × 1 vector of regression coefficients of the *m* genetic variants, and *ϵ_k_* is an *n* × 1 vector of non-systematic error term with each element following 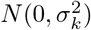. To test for *H_0_* : *β_.k_* = 0, a variance component test under the mixed effects model have been proposed to increase power over the usual F-test(Wu et al., 2011). The variance component test assumes that the regression coefficients, *β_.k_*, are random variables and follow a centered distribution with variance *τ*^2^Σ*_G_* (see below). Under these assumptions, the test for *β_.k_* = 0 is equivalent to testing *τ* = 0. The score statistic for this test is

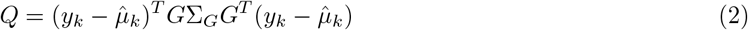

where 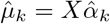 is the estimated mean of *y_k_* under the null hypothesis of no association. The test statistic Q asymptotically follows a mixture of chi-squared distributions under the null hypothesis and p-values can be computed by inverting the characteristic function (Davies, 1980).

The kernel matrix Σ*_G_* plays a critical role; it models the relationship among the effect sizes of the variants on the phenotypes. Any positive semidefinite matrix can be used for Σ*_G_* providing a unified framework for the region-based tests. A frequent choice of Σ*_G_* is a sandwich type matrix Σ*_G_* = *WR_G_W*, where *W* = *diag*(*w*_1_,.., *w_m_*) is a diagonal weighting matrix for each variant, and *R_G_* is a correlation matrix between the effect sizes of the variants. *R_G_* = *I_m×m_* implies uncorrelated effect sizes and corresponds to SKAT, and *R_G_* = 1_*m*_1_*m*_*^T^* corresponds to the burden test, where *I_m×m_* is an *m* × *m* diagonal matrix and 1_*m*_ = (1, ⋯ 1)*^T^* is an *m* × 1 vector with all elements being unity. Furthermore, a linear combination of these two matrices corresponds to *R_G_* = *ρ*1_*m*_1_*m*_*^T^* + (1 − *ρ*)*I_m×m_*, which is used for SKAT-O (Lee et al., 2012b).

### Multiple-phenotype region-based tests

Extending the idea of using kernels, we build a model for multiple phenotypes. The multivariate linear model shown in equation (3) can relate genetic variants to *K* correlated phenotypes,

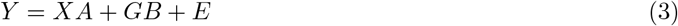

where *Y* = (*y*_1_, ⋯, *y_K_*) is an *n* × *K* phenotype matrix; *A* is a *q* × *K* matrix of coefficients of *X*; *B* = (*β_ij_*) is an *m* × *K* matrix of coefficients where *β_i_j* denotes the effect of the *i^th^* variant on the *j^th^* phenotype and *E* is an *n* × *K* matrix of non-systematic errors. Let *vec*(·) denote the matrix vectorization function, and then *vec*(*E*) follows *N*(0, *I_n_* ⊗ *V*), where *V* is a *K × K* covariance matrix and ⊗ represents the Kronecker product.

In addition to assuming that *β_.k_* follows a centered distribution with covariance *τ*^2^Σ*_G_*, we further assume that *β_i._* = (*β*_*i*1_, ⋯, *β_iK_*)*^T^*, which is the vector of regression coefficients of variant *i* for *K* multiple phenotypes, follows a centered distribution with covariance *τ*^2^Σ*_P_*, which implies that *vec*(*B*) follows a centered distribution with covariance *τ*^2^Σ*_G_* ⊗ Σ*_P_*. As before, the null hypothesis *H*_0_ : *vec*(*B*) = 0 is equivalent to *τ* = 0. The corresponding score test statistic is

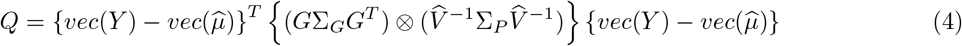

where 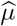 and 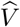 are the estimated mean and covariance of *Y* under the null hypothesis.

Σ*_P_* plays a similar role as Σ*_G_* but with respect to phenotypes. Σ*_P_* represents a kernel in the phenotypes space and models the relationship among the effect sizes of a variant on each of the phenotypes. Any positive semidefinite matrix can be used as Σ*_P_*.

The proposed approach provides a double-flexibility in modeling. Through the choice of structures for Σ*_G_* and *Σ_P_*, we can control the dependencies of genetic effects. Additionally, similar to SKAT, the use of a sandwich type matrix *WR_G_W* for Σ*_G_* allows us to upweight rare variants by using *Beta*(1, 25) weights as in Wu et al (Wu et al., 2011). Most of our hypotheses about the underlying genetic structure of a set of phenotypes can be modeled through varying structures of these two matrices.

### Phenotype kernel structure Σ*_P_*

The use of Σ*_G_* has been extensively studied previously in literature (Wu et al., 2011; Lee et al., 2012b,a). Here we propose several choices for Σ_*P*_ and study their effect from a modeling perspective.

#### Homogeneous (Hom) Kernel

It is possible that effect sizes of a variant on different phenotypes are homogeneous, in which case *β*_*j*1_ = ⋯ = *β_jK_*. Under this assumption,

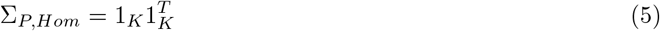

Under Σ*_P,Hom_*, the effect sizes *β_jk_*, (*k* = 1, ⋯ *K*) for a variant *j* are the same for all the phenotypes.

#### Heterogeneous (Het) Kernel

Effect sizes of a variant on different phenotypes can be heterogeneous in which *β*_*j*1_ ≠ ⋯ ≠ *β_jK_*. Under this assumption, we can construct

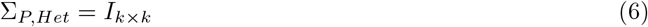

The *Σ_P,Het_* implies that the effect sizes 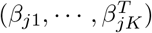 are uncorrelated among themselves. This also indicates that the correlation among the phenotypes is not affected by this particular region or gene.

#### Phenotype Covariance (PhC) Kernel

We may model Σ_*P*_ as proportional to the estimated residual covariance across the phenotypes as,

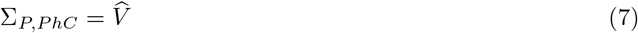

where 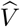 is the estimated covariance matrix among the phenotypes. This model assumes that the correlation between the effect sizes is proportional to that between the residual phenotypes after adjusting for the non-genetic covariates.

#### Principal Component (PC) Kernel

Principal component analysis (PCA) is a popular tool for multivariate analysis. In multiple phenotype tests, PC-based approaches have been used to reduce the dimension in phenotypes (Aschard et al., 2014). Here we show that PC-based approach can be included in our framework. Let *L* = (*L*_1_, ⋯, *L_K_*) be the loading matrix with each column *L_i_* produces the *i^th^* PC score. In Appendix A, we show that using 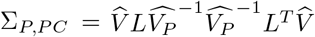 is equivalent to assuming heterogeneous effects with all PCs as phenotypes. Instead of using all the PC’s, we can use selected PC’s that represent the majority of cumulative variation in phenotypes. For example, we can jointly test the PC’s that have cumulative variance of 90%. If the top *t* PC’s have been chosen for analysis using *ν*% cumulative variance as cutoff, we can use

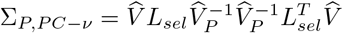

where *L_sel_* = [*L*_1_, ⋯, *L_t_*, 0, ⋯, 0] and 0 represents a vector of 0’s of appropriate length.

#### Relationship with other Multiple-Phenotype rare variant tests

We have proposed a uniform framework of Multi-SKAT tests that depend on Σ*_G_* and Σ*_P_*. There are certain specific choices of kernels that correspond to other published methods.

- Using Σ*_P,PhC_* and Σ*_G_* = *WI_m_W^T^* is identical to the GAMuT(Broadaway et al., 2016) with the projection phenotype kernel and the MSKAT with the *Q* statistic (Wu and Pankow, 2016).
- Using 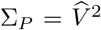 and Σ*_G_* = *WI_m_W^T^* is identical to GAMuT(Broadaway et al., 2016) with the linear phenotype kernel and the MSKAT with the *Q*′ statistic (Wu and Pankow, 2016).
- Using Σ*_P,Hom_* and Σ*_G_* = *WI_m_W^T^* is identical to hom-MAAUSS (Lee et al., 2016).
- Using Σ*_P,Het_* and Σ*_G_* = *WI_m_W^T^* is identical to het-MAAUSS (Lee et al., 2016) and MF-KM (Yan et al., 2015).

For the detailed proof, please see Appendix B.

### Minimum p-value based omnibus tests (minP & minP_com_)

The model and the corresponding test of association that we propose has two parameters, Σ_*G*_ and Σ_*P*_, which are absent in the null model of no association. Since our test is a score test, Σ_*G*_ and Σ_*P*_ cannot be estimated from the data. One possible approach is to select Σ*_G_* and Σ*_P_* based on prior knowledge; however, if the selected Σ_*G*_ and Σ_*P*_ do not reflect underlying biology, the test may have substantially reduced power (Ray et al., 2016; Lee et al., 2016). In an attempt to achieve robust power, we aggregate results across different Σ_*G*_ and Σ_*P*_ using the minimum of p-values from different kernels.

Although this omnibus test approach has been used in rare variant tests and multiple phenotype analysis for combining multiple kernels from genotypes and phenotypes (Zhan et al., 2017; Wu et al., 2013; Urrutia et al., 2015; He et al., 2017), it is challenging to calculate the p-value, since the minimum p-value does not follow the uniform distribution. One possible approach is using permutation or perturbation to calculate the monte-carlo p-value (Urrutia et al., 2015; Zhan et al., 2017); however, this approach is computationally too expensive to be used in genome-wide analysis. To address it, here we propose a fast copula based p-value calculation for Multi-SKAT, which needs only a small number of resampling steps to calculate the p-value.

Suppose *p_h_* is the p-value for *Q_h_* with given *h^th^* Σ*_G_* and Σ*_P_*, *h* = 1, ⋯, *b*, and *T_P_* = (*p*_1_, ⋯, *p_b_*)*^T^* is an *b* × 1 vector of p-values of *b* such Multi-SKAT tests. The minimum p-value test statistic after the Bonferroni adjustment is *b × p_min_*, where *p_min_* is the minimum of the *b* p-values. In the presence of positive correlation among the tests, this approach is conservative and hence might lack power of detection. Rather than using Bonferroni corrected *p_min_*, more accurate results can be obtained if the joint null distribution or more specifically the correlation structure of *T_P_* can be estimated. Here we adopt a resampling based approach to estimate this correlation structure. Note that our test statistic is equivalent to

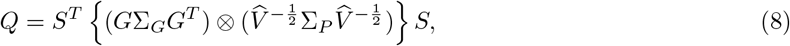

where 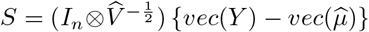. Under the null hypothesis *S* approximately follows an uncorrelated normal distribution *N*(0, *I_nK_*). Using this, we propose the following resampling algorithm

- Step 1. Generate *nK* samples from an *N*(0, 1) distribution, say *S_R_*.
- Step 2. Calculate *b* different test statistics as 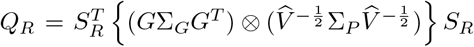 for all the choices of Σ_*P*_ and calculate p-values.
- Step 3. Repeat the previous steps independently for *R*(= 1000) iterations, and calculate the correlation between the p-values of the tests from the *R* resampling p-values.

With the estimated null correlation structure, we use a Copula to approximate the joint distribution of *T_P_* (Demarta and McNeil, 2005; He et al., 2017). Copula is a statistical approach to construct joint multivariate distribution using marginal distribution of each variable and correlation structure. Since marginally each test statistic *Q* follows a mixture of chi-square distributions, which has a heavier tail than normal distribution, we propose to use a t-Copula to approximate the joint distribution, i.e, we assume the joint distribution of *T_P_* to be multivariate t with the estimated correlation structure. The final p-value for association is then calculated from the distribution function of the assumed t-Copula.

When calculating the correlation across the p-values, Pearson’s correlation coefficient can be unreliable since it depends on normality and homoscedasticity assumptions. To avoid such assumptions we recommend estimating the null-correlation matrix of the p-values through Kendall’s tau (*τ*), which is a non-parametric approach based on concordance of ranks.

The minimum p-value approach can be used to combine different Σ*_P_* given Σ*_G_*, or combine both Σ*_P_* and Σ*_G_*. For example two Σ*_G_*’s corresponding to SKAT (*WW*) and Burden kernels 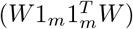 and four Σ*_P_*’s (Σ*_P,Hom_*, Σ*_P,Het_*, *Σ_P,PhC_*, Σ_*P,PC*−0.9_) can be comnibed, which results in the omnibus test of these eight different tests. To differentiate the latter, we will call it minP_com_ which combines SKAT and Burden type kernels of Σ*_G_*.

### Adjusting for relatedness

We formulated equation (3) and corresponding tests under the assumption of independent individuals. If individuals are related, this assumption is no longer valid, and the tests may have inflated type I error rate. Since our method is regression-based, we can relax the independence assumption by introducing a random effect term to account for the relatedness among individuals.

Let Φ be the kinship matrix of the individuals and *V_g_* is a co-heritability matrix, denoting the shared heritability between the phenotypes. Extending the model presented in equation (3), we incorporate Φ and *V_g_* as

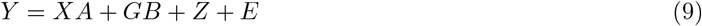

where *Z* is an *n × K* matrix with *vec*(*Z*) following *N*(0, Φ ⊗ *V_g_*). *Z* represents a matrix of random effects arising from shared genetic effects between individuals due to the relatedness. The remaining terms are the same as in equation (3). The corresponding score test statistic is

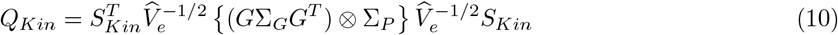

where 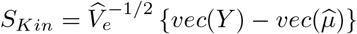 and 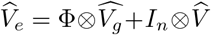 is the estimated covariance matrix of *vec*(*Y*) under the null hypothesis. Similar to the previous versions for unrelated individuals, *Q_Kin_* asymptotically follows a mixture of chi-square under the assumption of no association.

This approach depends on the estimation of the matrices Φ, *V_g_* and *V*. The kinship matrix Φ can be estimated using the genome-wide genotype data (Manichaikul et al., 2010). Several of the published methods like LD-Score(Bulik-Sullivan et al., 2015), PHENIX(Dahl et al., 2016) and GEMMA(Zhou and Stephens, 2014; Zhou et al., 2013) can jointly estimate *V_g_* and *V*. In our numerical analysis, we have used PHENIX. This is an efficient method to fit local maximum likelihood variance components in a multiple phenotype mixed model through an E-M algorithm.

Once the matrices Φ, *V_g_* and *V* are estimated, we compute the asymptotic p-values for *Q_Kin_* by using a mixture of chi-square distribution. The computation of *Q_Kin_* requires large matrix multiplications, which can be time and memory consuming. To reduce computational burden, we employ several transformations. We perform an eigen-decomposition on the kinship matrix Φ as Φ = *U*Λ*U^T^*, where *U* is an orthogonal matrix of eigenvectors and Λ is a diagonal matrix of corresponding eigenvalues. We obtain the transformed phenotype matrix as *Ỹ* = *YU*, the transformed covariate matrix as 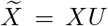, the transformed random effects matrix 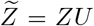 and transformed residual error matrix *Ẽ* = *EU*. Equation (9) can be transformed into

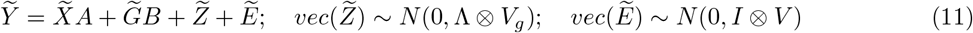

All the properties of the tests developed from equation (3) are directly applicable to those from equation 11). *Q_Kin_* can be computed from this transformed equation as,

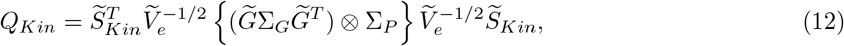

where 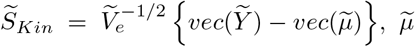 is the estimated mean of *Ỹ* under the null hypothesis and 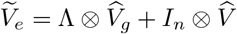. Asymptotic p-values can be obtained from the corresponding mixture of chi-squares distribution. Further, omnibus strategies for the tests developed from equation (3) are applicable in this case with similar modifications. For example, the resampling algorithm for minimum p-value based omnibus test can be implemented here as well by noting that 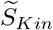 approximately follows an uncorrelated normal distribution.

### Simulations

We carried out extensive simulation studies to evaluate the type I error control and power of Multi-SKAT tests. For type I error simulations without related individuals and all power simulations, we generated 10,000 chromosomes over 1Mbp regions using a coalescent simulator with European demographic model (Schaffner et al., 2005). The MAF spectrum of the simulated variants is shown in Supplementary Figure S6, showing that most of the variants are rare variants. Since the average length of the gene is 3 kbps we randomly selected a 3 kbps region for each simulated dataset to test for associations. For the type I error simulations with related individuals, to have a realistic kinship structure, we used the METSIM study genotype data.

Phenotypes were generated from the multivariate normal distribution as

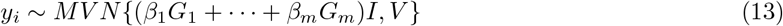

where *y_i_* = (*y*_*i*1_, ⋯, *y_iK_*)*^T^* is the outcome vector, *G_j_* is the genotype of the *j^th^* variant, and *β_j_* is the corresponding effect size, and *V* is a covariance of the non-systematic error term. We use *V* to define level of covariance between the traits. *I* is a *k* × 1 indicator vector, which has 1 when the corresponding phenotype is associated with the region and 0 otherwise. For example, if there are 5 phenotypes and the last three are associated with the region, *I* = (0, 0, 1, 1, 1) *^T^*.

To evaluate whether Multi-SKAT can control type I error under realistic scenarios, we simulated a dataset with 9 phenotypes with a correlation structure identical to that of 9 amino acid phenotypes in the METSIM data (See Supplementary Figure S1). Phenotypes were generated using equation (13) with *β* = 0. Total 5, 000, 000 datasets with 5, 000 individuals were generated to obtain the empirical type-I error rates at *α* = 10^−4^,10^−5^ and 2.5 × 10^−6^, which are corresponding to candidate gene studies to test for 500 and 5000 genes and exome-wide studies to test for all 20, 000 protein coding genes, respectively.

Next, we evaluated type I error controls in the presence of related individuals. To have a realistic related structure we used the METSIM study genotype data. We generated a random subsample of 5000 individuals from the METSIM study individuals and generated null values for the 9 phenotypes from *MVN*(0, *V_e_*), where 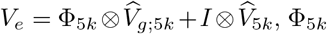 is the estimated kinship matrix of the 5000 selected individuals, 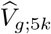 and 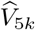 are estimated co-heritability and residual variance matrices respectively for these individuals as estimated using the MPMM function in the PHENIX R-package (version 1.0). For each set of 9 phenotypes, we performed the Multi-SKAT tests for a randomly selected 5000 genes in the METSIM data. For the details about the data, see next section. We carried out this procedure 1000 times and obtained 5, 000, 000 p-values, and estimated type I error rate as proportions of p-values smaller than the given level *α*.

Our simulation studies focus on evaluating the power of the proposed tests when the number of phenotypes are 5 or 6. Power simulations were performed both in situations when there was no pleiotropy (i.e., only one of the phenotypes was associated with the causal variants) and also when there was pleiotropy. Under pleiotropy, since it is unlikely that all the phenotypes are associated with genotypes in the region, we varied the number of phenotypes associated. For each associated phenotype, 30% or 50% of the rare variants (MAF < 1%) were randomly selected to be causal variants. We modeled the rarer variants to have stronger effect, as |*β_j_*| = *c*|*log*_10_(*MAF_j_*)|. We used *c* = 0.3 which yields |*β_j_*| = 0.9 for variants with MAF = 10^−3^. Our choice of *β* yielded the average heritability of associated phenotypes between 1% to 4%. We also considered situations that all causal variants were trait-increasing variants (i.e. positive β) or 20 % of causal variants were trait-decreasing variants (i.e. negative β). Empirical power was estimated from 1000 independent datasets at exome-wide *α* = 2.5 × 10^−6^.

In type I error and power simulations, we compared the following tests:

- Bonferroni adjusted minimum p-values from gene-based test (SKAT, Burden or SKAT-O) on each phenotype (minPhen)
- Multi-SKAT with Σ*_P,Hom_* (Hom)
- Multi-SKAT with Σ*_P,Het_* (Het)
- Multi-SKAT with Σ*_P,PhC_* (PhC)
- Multi-SKAT with Σ_*P,PC*−0.9_ (PC-Sel)
- Minimum P-value of Hom, Het, PhC and PC-Sel using Copula (minP)
- Minimum P-value of Hom, Het, PhC and PC-Sel with Σ*_G_* being SKAT and Burden, using Copula (minP_com_)

For the Multi-SKAT tests, we used two different Σ*_G_*’s corresponding to SKAT (i.e. Σ*_G_* = *WW*) and Burden tests (i.e. 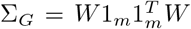). For the variant weighting matrix *W* = *diag*(*w*_1_, ⋯, *w_m_*), we used *w_j_* = *Beta*(*MAF_j_*, 1, 25) function to upweight rarer variants, as recommended by Wu et al. (Wu et al., 2011).

### Computation Time

We estimated the computation time of Multi-SKAT tests and the existing methods. Using simulated datasets of 5000 related and unrelated individuals with 10 phenotypes and 20 genetic variants, we estimated the computation time of Multi-SKAT tests with and without kinship adjustments. To compare the computation performance of Multi-SKAT tests with the existing methods, we generated datasets of unrelated individuals with five different sample sizes (n = 1000, 2000, 5000, 10000, 15000 and 20000) and four different number of variants (*m* = 10, 20, 50,100). For each simulation setup, we generated 100 datasets and obtained the average value of the computation time.

### Analysis of the METSIM study exomechip data

To investigate the cross-phenotype roles of low frequency and rare variants on amino acids, we analyzed data on 8545 participants of the METSIM study on whom all 9 amino acids (Alanine, Leucine, Isoleucine, Glycine, Valine, Tyrosine, Phenylalanine, Glutamine, Histidine) were measured by proton nuclear magnetic resonance spectroscopy(Teslovich et al., 2018). Individuals were genotyped on the Illumina ExomeChip and OmniExpress arrays and we included individuals that passed sample QC filters (Huyghe et al., 2013). The kinship between the individuals was estimated via KING (version 2.0) (Manichaikul et al., 2010). We adjusted the amino acid levels for age, age^2^ and BMI and inverse-normalized the residuals. The phenotype correlation matrix after covariate adjustment is shown in Figure 3 and Supplementary Figure S1. Subsequently, we estimated the genetic heritability matrix and the residual covariance matrix using the MPMM function from PHENIX (Dahl et al., 2016) R package.

We included rare (MAF < 1%) nonsynonymous and protein-truncating variants with a total rare minor allele count of at least 5 for genes that had at least 3 rare variants leaving 5207 genes for analysis. We set a stringent significance threshold at 9.6 × 10^−6^ corresponding to the Bonferroni adjustment for 5207 genes. Further, we also considered a less stringent threshold of 10^−4^, corresponding to a candidate gene study of 500 genes, as suggestive to study the associations which were not significant but close to the threshold.

## Results

### Type I Error simulations

We estimated empirical type I error rates of the Multi-SKAT tests with and without related individuals. For unrelated individuals, we simulated 5, 000 individuals and 9 phenotypes based on the correlation structure for the amino acids phenotypes in the METSIM study data. For related individuals, we simulated 5, 000 individuals using the kinship matrix for randomly chosen METSIM individuals (see the Method section). We performed association tests and estimated type I error rate as the proportion of p-values less than the specified *α* levels. Type I error rates of the Multi-SKAT tests were well maintained at *α* = 10^−4^, 10^−5^ and 2.5 × 10^−6^ for both unrelated and related individuals (Table 1), which correspond to candidate gene studies of 500 and 5000 genes and exome-wide studies to test for all 20, 000 protein coding genes, respectively. For example, at level *α* = 2.5 × 10^−6^, the largest empirical type I error rate from any of the Multi-SKAT tests was 3.4 × 10^−6^, which was within the 95% confidence interval (CI = (1.6 × 10^−6^, 4 × 10^−6^)).

**Table 1:** Empirical type I error rates of the Multi-SKAT tests from 5,000,000 simulated null datasets. The number of phenotypes were nine and the correlation structure among the phenotypes were similar to that of the amino acid phenotypes in the METSIM study data. The sample size was 5000.

| | Level | $\Sigma_G$ | minPhen | Hom | Het | PhC | PC-Sel | minP | minP<sub>com</sub> |
| --- | --- | --- | --- | --- | --- | --- | --- | --- | --- |
| Independent samples<br>(without kinship adjustment) | $2.5 \times 10^{-6}$ | *SKAT* | $2.4 \times 10^{-06}$ | $2.6 \times 10^{-06}$ | $2.6 \times 10^{-06}$ | $2.8 \times 10^{-06}$ | $2.6 \times 10^{-06}$ | $3.4 \times 10^{-06}$ | $2.8 \times 10^{-06}$ |
| | | *Burden* | $2.4 \times 10^{-06}$ | $2.8 \times 10^{-06}$ | $2.6 \times 10^{-06}$ | $2.6 \times 10^{-06}$ | $2.6 \times 10^{-06}$ | $3.0 \times 10^{-06}$ | |
| | $10^{-05}$ | *SKAT* | $9.6 \times 10^{-06}$ | $9.6 \times 10^{-06}$ | $9.8 \times 10^{-06}$ | $9.8 \times 10^{-06}$ | $9.8 \times 10^{-06}$ | $9.4 \times 10^{-06}$ | $1.2 \times 10^{-05}$ |
| | | *Burden* | $9.4 \times 10^{-06}$ | $9.6 \times 10^{-06}$ | $9.8 \times 10^{-06}$ | $9.6 \times 10^{-06}$ | $9.6 \times 10^{-06}$ | $9.6 \times 10^{-06}$ | |
| | $10^{-04}$ | *SKAT* | $9.6 \times 10^{-05}$ | $9.4 \times 10^{-05}$ | $9.6 \times 10^{-05}$ | $9.8 \times 10^{-05}$ | $9.8 \times 10^{-05}$ | $9.8 \times 10^{-05}$ | $1.1 \times 10^{-04}$ |
| | | *Burden* | $9.8 \times 10^{-05}$ | $9.6 \times 10^{-05}$ | $9.4 \times 10^{-05}$ | $9.7 \times 10^{-05}$ | $9.6 \times 10^{-05}$ | $9.7 \times 10^{-05}$ | |
| Related samples<br>(with kinship adjustment) | $2.5 \times 10^{-6}$ | *SKAT* | $2.2 \times 10^{-06}$ | $2.4 \times 10^{-06}$ | $2.4 \times 10^{-06}$ | $2.8 \times 10^{-06}$ | $2.6 \times 10^{-06}$ | $3.0 \times 10^{-06}$ | $2.6 \times 10^{-06}$ |
| | | *Burden* | $2.2 \times 10^{-06}$ | $2.6 \times 10^{-06}$ | $2.4 \times 10^{-06}$ | $2.6 \times 10^{-06}$ | $2.4 \times 10^{-06}$ | $3.2 \times 10^{-06}$ | |
| | $10^{-05}$ | *SKAT* | $9.6 \times 10^{-06}$ | $9.8 \times 10^{-06}$ | $9.8 \times 10^{-06}$ | $9.8 \times 10^{-06}$ | $9.8 \times 10^{-06}$ | $9.4 \times 10^{-06}$ | $1.4 \times 10^{-05}$ |
| | | *Burden* | $9.7 \times 10^{-06}$ | $9.6 \times 10^{-06}$ | $9.4 \times 10^{-06}$ | $9.4 \times 10^{-06}$ | $9.4 \times 10^{-06}$ | $9.4 \times 10^{-06}$ | |
| | $10^{-04}$ | *SKAT* | $9.4 \times 10^{-05}$ | $9.6 \times 10^{-05}$ | $9.7 \times 10^{-05}$ | $9.8 \times 10^{-05}$ | $9.4 \times 10^{-05}$ | $9.8 \times 10^{-05}$ | $1.2 \times 10^{-04}$ |
| | | *Burden* | $9.5 \times 10^{-05}$ | $9.7 \times 10^{-05}$ | $9.6 \times 10^{-05}$ | $9.8 \times 10^{-05}$ | $9.8 \times 10^{-05}$ | $9.7 \times 10^{-05}$ | |

### Power simulations

We compared the empirical power of the minPhen (Bonferroni adjusted minimum p-value for the phenotypes) and Multi-SKAT tests. For each simulation setting, we generated 1,000 sequence datasets of 5, 000 unrelated individuals and for each test estimated empirical power as the proportion of p-values less than *α* = 2.5 × 10^−6^, reflecting Bonferroni correction for testing 20,000 independent genes. Since the Hom and Het tests are identical to hom-MAAUSS and het-MAAUSS, respectively, and using PhC is identical to both GAMuT (with projection phenotype kernel) and MSKAT, our power simulation studies effectively compare the existing multiple phenotype tests.

In Figure 1, we show the results for 5 phenotypes with compound symmetric correlation structure with the correlation 0.3 or 0.7, where 30% of rare variants (MAF < 0.01) were positively associated with 1, 2 or 3 phenotypes. Since it is unlikely that all the phenotypes are associated with the region, we restricted the number of associated phenotypes to at most 3. In most scenarios, PhC, PC-Sel and Het had greater power among the Multi-SKAT tests with fixed phenotype kernels (i.e. Hom, Het, PhC and PC-Sel) while minP, maintained high power as well. For example, when the correlation between the phenotypes was 0.3 (i.e. *ρ* = 0.3) and SKAT kernel was used for the genotype kernel Σ*_G_*, if 3 phenotypes were associated with the region, minP and PhC were more powerful than the other tests. If the correlation between the phenotypes was *ρ* = 0.7 and Burden kernel was used for genotype kernel Σ*_G_*, Het, PC-Sel and minP had higher power than the rest of the tests when 2 phenotypes were associated. It is noteworthy that Hom had the lowest power in all the scenarios of Figure 1.

**Figure 1:**
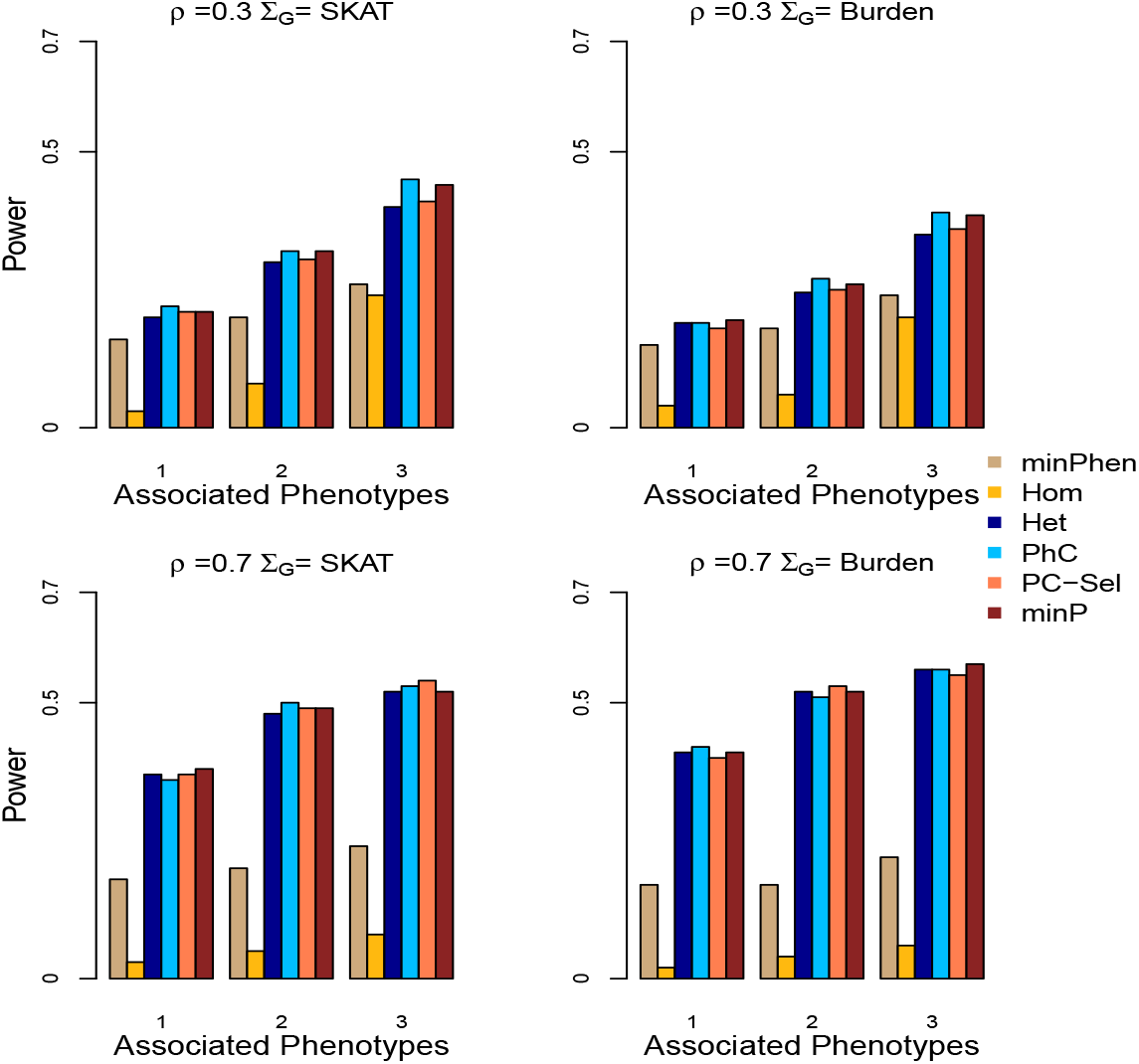
Power for Multi-SKAT tests when phenotypes have compound symmetric correlation structures. Empirical power for minPhen, Hom, Het, PhC, PC-Sel, minP plotted against the number of phenotypes associated with the gene of interest with a total of 5 phenotypes under consideration. Upper row shows the results for *ρ* = 0.3 and lower row for *ρ* = 0.7. Left column shows results with SKAT kernel Σ*_G_*, and right columns shows results with Burden kernel. All the causal variants were trait-increasing variants.

Figure 2 demonstrates scenarios involving 6 phenotypes and clustered correlation structures where PhC was outperformed by other choices of the phenotype kernel Σ*_P_*. When all three phenotype clusters had associated phenotypes and the correlation within the clusters was low (*ρ* = 0.3) (Figure 2, upper panel), Hom and minP tests outperformed PhC when the SKAT kernel was used. This may be because that the phenotype correlation structure did not reflect the genetic association pattern. When 2 small clusters had high within-cluster correlation (*ρ* = 0.7) and one large cluster had low within-cluster correlation (*ρ* = 0.3) (Figure 2, lower panel), Het and minP had higher power than PhC.

**Figure 2:**
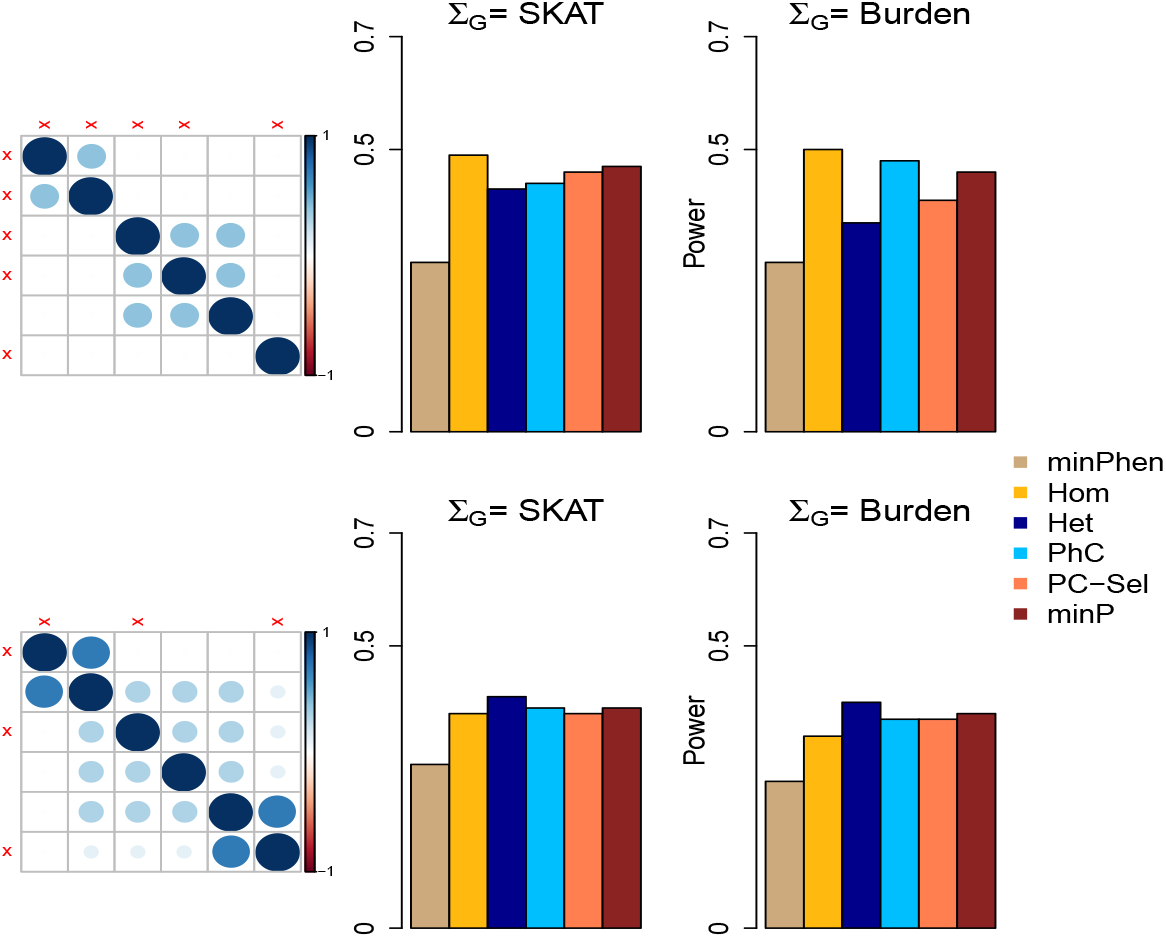
Power for Multi-SKAT tests when phenotypes have clustered correlation structures. Empirical powers for minPhen, Hom, Het, PhC, PC-Sel, minP are plotted under different levels of association with a total of 6 phenotypes and with clustered correlation structures. Middle column shows the empirical powers for different combinations of phenotypes associated with SKAT kernel Σ*_G_*; the rightmost column shows the corresponding results with Burden kernel; left column shows the corresponding correlation matrices for the phenotypes. The associated phenotypes are indicated in red cross marks across the correlation matrices. All the causal variants were trait-increasing variants.

When 20% of causal variants were trait-decreasing variants (80% trait-increasing), the power of Multi-SKAT tests with Burden Σ*_G_* was reduced (Supplementary Figure S2 and S3). This is because the association signals were attenuated due to the mix of trait-increasing and trait-decreasing variants. Since SKAT is robust regardless of the association direction, power with SKAT Σ*_G_* was largely maintained. The relative performance of methods with different Σ*_P_* given Σ*_G_* was quantitatively similar to the results without trait-decreasing variants.

Further, we estimated power of minP_com_, which combines tests across phenotype (Σ*_P_*) and genotype Σ*_G_* kernels. The power of minP_com_ was evaluated for the compound symmetric phenotype correlation structure presented in Figure 1 and was compared with the two minP tests of SKAT (minP-SKAT) and Burden (minP-Burden) Σ*_G_* kernels. Figure 3 shows empirical power with and without trait-decreasing variants. When all genetic effect coefficients were positive (Figure 3, left panel) the performances of minP-SKAT and minP-Burden were similar for both the situations where the correlation between the phenotypes were low (i.e. *ρ* = 0.3) and high (i.e. *ρ* = 0.7). When 20% of genetic effect coefficients were negative (Figure 3, right panel), as expected, the power of minP-Burden was substantially decreased. Across all the situations, the power of minP_com_ was similar to the most powerful minP with fixed genotype kernel Σ*_G_*. When 50% of variants were causal variants and all genetic effect coefficients were positive (Supplementary Figure S4, left panel), minP-Burden was more powerful than minP-SKAT, and minP_com_ had similar power than minP-Burden.

**Figure 3:**
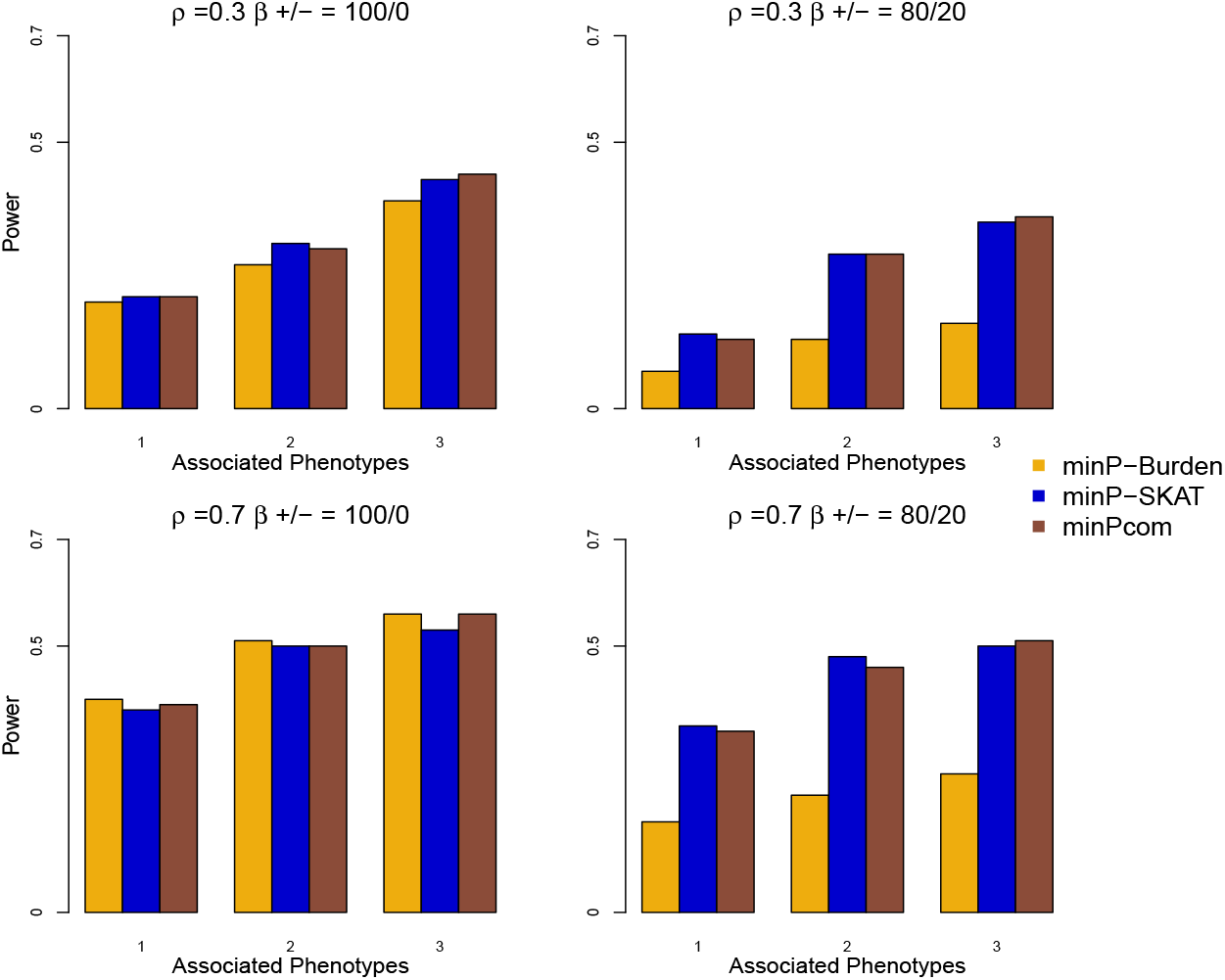
Power for Multi-SKAT by combining tests with Σ*_P_* as Hom, Het, PhC, PC-Sel and Σ*_G_* as SKAT and Burden when phenotypes have compound symmetric correlation structures. Empirical powers for minP-Burden, minP-SKAT and minP_com_ are plotted against the number of phenotypes associated with the gene of interest with a total of 5 phenotypes under consideration. Upper row shows the results for *ρ* = 0.3 and lower row for *ρ* = 0.7. Left column shows results when all the causal variants were trait-increasing variants, and right column shows results when 80%/20% of the causal variants were trait-increasing/trait-decreasing variants.

Overall, our simulation results show that the omnibus tests, especially minP_com_, had robust power throughout all the simulation scenarios considered. When Σ*_G_* and Σ*_P_* were fixed, power depended on the model of association and the correlation structure of the phenotypes. Overall, the proposed Multi-SKAT tests generally outperformed the single phenotype test (minPhen), even when only one phenotype was associated with genetic variants.

### Application to the METSIM study exomechip data

Inborn errors of amino acid metabolism cause mild to severe symptoms including type 2 diabetes(Stančáková et al., 2012; Würtz et al., 2012, 2013) and liver diseases(Tajiri and Shimizu, 2013) among others. Amino acid levels are perturbed in certain disease states, e.g., glutamic and aspartic acid levels are reduced in Alzheimer disease brains(Allan Butterfield and Pocernich, 2003); Isoleucine, glycine, alanine, phenylalanine, and threonine levels are increased in cerebo-spinal fluid (CSF) of individuals with motor neuron disease(de Belleroche et al., 2003). To find rare variants associated with the 9 measured amino acid levels, we applied the MultiSKAT tests to the METSIM study data(Teslovich et al., 2018). The MAF spectrum of the genotyped variants is shown in Supplementary Figure S6, showing that most of the variants are rare variants. We estimated the relatedness between individuals by KING (Manichaikul et al., 2010), and coheritability of the amino acid phenotypes and the corresponding residual variance using PHENIX(Dahl et al., 2016) (Supplementary Figure S1). Among the 8,545 METSIM participants with non-missing phenotypes and covariates, 1,332 individuals had a second degree or closer relationship with one or more of the METSIM participants. A total of 5, 207 genes with at least three rare variants were included in our analysis. The Bonferroni corrected significance threshold was *α*= 0.05/5207 = 9.6 × 10^−6^. Further we used a less significant cutoff of *α*= 10^−4^ for a gene to be suggestive. After identifying associated genes, we carried out backward elimination procedure (Appendix C) to investigate which phenotypes are associated with the gene. This procedure iteratively removes phenotypes based on minP_com_ p-values.

QQ plots for the p-values obtained by minPhen and Multi-SKAT omnibus tests (minP and minP_com_) are displayed in Figure 4. Due to the presence of several strong associations, for the ease of viewing, any p-value < 10^−12^ was collapsed to 10^−12^. The QQ plots are well calibrated with slight inflation in tail areas. The genomic-control lambda (*λ_GC_*) varied between 0.97 and 1.04, which indicates no inflation of test statistics. Table 2 shows genes with p-values less than 10^−4^ for minPhen or minP_com_. Table 5 shows SKAT-O p-values for each of the gene - amino acid pairs. Among the eight significant or suggestive genes displayed in Table 2, minP_com_ provides more significant p-values than minPhen for six genes: Glycine decarboxylase (*GLDC* [MIM: 238300]), Histidine ammonia-lyase (*HAL* [MIM: 609457]), Phenylalanine hydroxylase (*PAH* [MIM: 612349]), Dihydroorotate dehydrogenase (*DHODH* [MIM: 126064]), Mediator of RNA polymerase II transcription subunit 1 (*MED1* [MIM: 604311]), Serine/Threonine Kinase 33 (*STK33* [MIM: 607670]). Interestingly, *PAH* and *MED1* are significant by minP_com_, but not significant by minPhen. *PAH* encodes an Phenylalanine hydroxylase, which catalyzes the hydroxylation of the aromatic side-chain of phenylalanine to generate tyrosine. *MED1* is involved in the regulated transcription of nearly all RNA polymerase II-dependent genes. This gene does not show any single phenotype association, but cross-phenotype analysis produced evidence of association. Using backward elimination we find that Phenylalanine and Tyrosine are the last two phenotypes to be eliminated (Supplementary Table S2). We have provided a detailed description of the function and clinical implications of the significant and suggestive genes in Supplementary Table S4.

**Figure 4:**
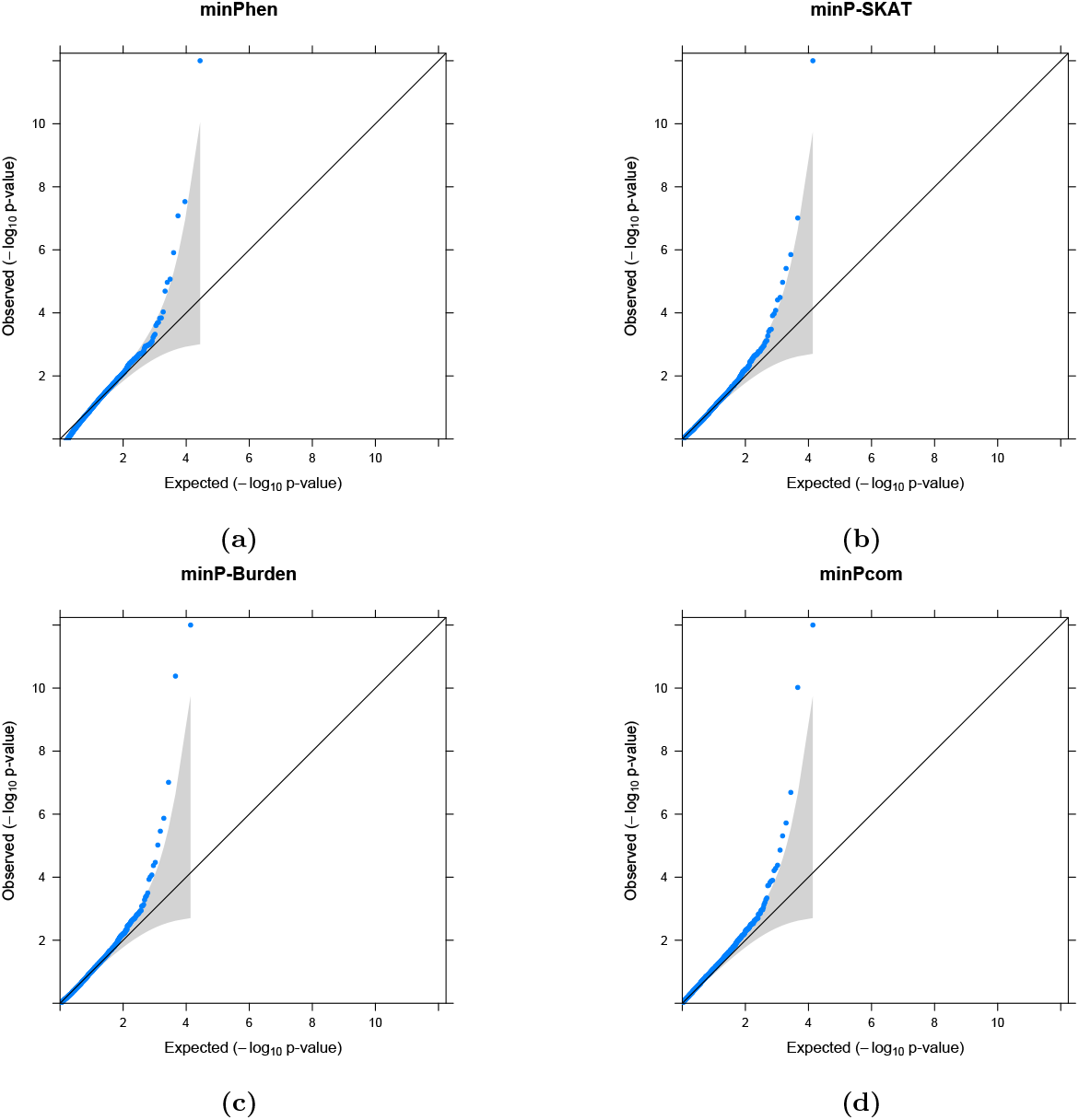
QQplot of the p-values of minPhen and Multi-SKAT omnibus tests for the METSIM data. For the ease of viewing, any associations with p-values < 10^−12^ have been collapsed to 10^−12^.

**Table 2:** Significant and suggestive genes associated with 9 amino acid phenotypes. Genes with p-values < 10^−4^ by minPhen or any Multi-SKAT tests (minP-SKAT, minP-Burden and minP_com_) were reported in this table. Multi-SKAT tests were applied with the kinship adjustment, and 5207 genes with at least three rare (MAF < 0.01) nonsynonymous and protein-truncating variants were used in this analysis. The total sample size was n=8545. P-values smaller than the Bonferroni corrected significance *α* = 9.6 × 10^−6^ were marked as bold. Smallest p-value for each gene among all the tests have been underlined. minPhen was calculated as the Bonferroni adjusted minimum SKAT-O p-value across each phenotype.

| Gene | Chromosome | Rare SNPs | MAC | minPhen | minP-SKAT | minP-Burden | minP<sub>com</sub> |
| --- | --- | --- | --- | --- | --- | --- | --- |
| *GLDC* | 9 | 4 | 183 | **$2.2 \times 10^{-63}$** | **$1.1 \times 10^{-72}$** | **$9.7 \times 10^{-64}$** | **$2.3 \times 10^{-72}$** |
| *HAL* | 3 | 6 | 42 | **$2.9 \times 10^{-08}$** | $1.1 \times 10^{-05}$ | **$4.2 \times 10^{-11}$** | **$9.5 \times 10^{-11}$** |
| *DHODH* | 16 | 5 | 90 | **$1.3 \times 10^{-06}$** | **$9.7 \times 10^{-08}$** | **$7.7 \times 10^{-06}$** | **$1.1 \times 10^{-07}$** |
| *PAH* | 12 | 6 | 27 | $6.1 \times 10^{-04}$ | **$1.4 \times 10^{-06}$** | **$1.3 \times 10^{-06}$** | **$1.9 \times 10^{-06}$** |
| *MED1* | 17 | 3 | 147 | $6.2 \times 10^{-01}$ | **$3.9 \times 10^{-06}$** | $3.4 \times 10^{-04}$ | **$4.9 \times 10^{-06}$** |
| *STK33* | 11 | 6 | 180 | $8.9 \times 10^{-01}$ | $3.2 \times 10^{-05}$ | $3.3 \times 10^{-02}$ | $4.2 \times 10^{-05}$ |
| *ALDH1L1* | 3 | 8 | 103 | **$8.4 \times 10^{-08}$** | $3.8 \times 10^{-04}$ | $4.2 \times 10^{-05}$ | $5.4 \times 10^{-05}$ |
| *BCAT2* | 19 | 3 | 133 | <u>$1.1 \times 10^{-05}$</u> | $9.3 \times 10^{-03}$ | $3.4 \times 10^{-05}$ | $6.2 \times 10^{-05}$ |

Among other genes, *GLDC* has the smallest p-value. Variants in *GLDC* are known to cause glycine encephalopathy (MIM: 605899) (Hughes, 2009). To investigate whether our results were supported by single phenotype associations, we applied SKAT-O to each of the 9 amino acid phenotypes. Univariate SKAT-O test with each of these phenotype reveals that this gene has a strong association with Glycine (p-value = 2.5 × 10^−64^, Table 5). Among the variants genotyped in this gene, rs138640017 (MAF = 0.009) appears to drive the association (single variant p-value =1.0 × 10^−64^). Variants in *HAL* cause histidinemia (MIM: 235800) in human and mouse. This gene shows significant univariate association with Histidine (SKAT-O p-value = 3.2 × 10^−8^, Table 5) which in turn is influenced by the association of rs141635447 (MAF = 0.005) with Histidine (single variant p-value = 3.7 × 10^−13^). Similarly, variants in *DHODH*, which have been previously found to be associated with postaxial acrofacial dysostosis (MIM: 263750), have significant cross-phenotype association although the result us mostly driven by the association with Alanine (SKAT-O p-value =1.4 × 10^−07^, Table 5). *ALDH1L1* catalyzes conversion of 10-formyltetrahydrofolate to tetrahydrofolate. Published results show that common variant rs1107366, 5kb upstream of *ALDH1L1*, is associated with Glycine-Seratinine ratio (Xie et al., 2013). Down-regulation of *BCAT2* in mice causes elevated serum branched chain amino acid levels and features of maple syrup urine disease.

Table 3 shows p-values of Multi-SKAT kernel and minP with two genotype kernels (SKAT and Burden). Among phenotype kernels, PhC and Het generally produced the smallest p-values. We further applied Multi-SKAT tests without kinship adjustment on the whole METSIM study individuals. As expected, this produced inflation in QQ plots (Supplementary Figure S5).

**Table 3:** P-values for MultiSKAT tests (Hom, Het, PhC, PC-Sel, minP) with SKAT and burden kernels for the genes reported in Table 2. Multi-SKAT tests were applied with the kinship adjustment, and 5207 genes with at least three rare (MAF < 0.01) nonsynonymous and protein-truncating variants were used in this analysis. The total sample size was n=8545. P-values smaller than the Bonferroni corrected significance *α* = 9.6 × 10^−6^ were marked as bold. For the upper part of the table, minPhen was calculated as the Bonferroni adjusted minimum SKAT p-value across each phenotype, while for the lower part it was calculated as the Bonferroni adjusted minimum Burden p-value across each phenotype.

|  |  | Gene |  |  |  | Multi-SKAT |  |  |  |  |
| --- | --- | --- | --- | --- | --- | --- | --- | --- | --- | --- |
| $\Sigma_G$ | Name | Chromosome | Rare SNPs | MAC | minPhen | Hom | Het | PhC | PC-Sel | minP |
| SKAT | *GLDC* | 9 | 4 | 183 | **$6.8 \times 10^{-63}$** | $1.3 \times 10^{-03}$ | **$1.3 \times 10^{-17}$** | **$2.9 \times 10^{-73}$** | **$5.3 \times 10^{-59}$** | **$1.1 \times 10^{-72}$** |
| | *DHODH* | 16 | 5 | 90 | **$5.1 \times 10^{-08}$** | $1.4 \times 10^{-01}$ | $3.2 \times 10^{-03}$ | **$3.1 \times 10^{-08}$** | **$7.1 \times 10^{-06}$** | **$9.7 \times 10^{-08}$** |
| | *PAH* | 12 | 6 | 27 | $4.3 \times 10^{-05}$ | $7.9 \times 10^{-02}$ | $1.8 \times 10^{-05}$ | **$3.9 \times 10^{-07}$** | $6.3 \times 10^{-04}$ | **$1.4 \times 10^{-06}$** |
| | *MED1* | 17 | 3 | 147 | $8.7 \times 10^{-02}$ | $3.4 \times 10^{-01}$ | **$9.3 \times 10^{-07}$** | $2.8 \times 10^{-04}$ | $1.9 \times 10^{-04}$ | **$3.9 \times 10^{-06}$** |
| | *HAL* | 12 | 6 | 42 | $3.8 \times 10^{-05}$ | $5.5 \times 10^{-01}$ | $3.5 \times 10^{-03}$ | **$2.9 \times 10^{-06}$** | $1.5 \times 10^{-04}$ | $1.1 \times 10^{-05}$ |
| | *STK33* | 11 | 6 | 180 | $4.0 \times 10^{-01}$ | $2.3 \times 10^{-01}$ | **$8.9 \times 10^{-06}$** | $1.1 \times 10^{-03}$ | $5.7 \times 10^{-03}$ | $3.2 \times 10^{-05}$ |
| | *ALDH1L1* | 3 | 8 | 103 | $4.4 \times 10^{-02}$ | $1.1 \times 10^{-01}$ | $2.2 \times 10^{-02}$ | $1.6 \times 10^{-04}$ | $4.2 \times 10^{-04}$ | $3.8 \times 10^{-04}$ |
| | *BCAT2* | 19 | 3 | 133 | $9.7 \times 10^{-04}$ | $5.0 \times 10^{-01}$ | $8.1 \times 10^{-02}$ | $2.4 \times 10^{-03}$ | $3.5 \times 10^{-03}$ | $9.3 \times 10^{-03}$ |
| Burden | *GLDC* | 9 | 4 | 183 | **$1.4 \times 10^{-59}$** | $7.5 \times 10^{-04}$ | **$1.5 \times 10^{-15}$** | **$1.3 \times 10^{-64}$** | **$2.1 \times 10^{-54}$** | **$9.7 \times 10^{-64}$** |
| | *HAL* | 12 | 6 | 42 | **$2.4 \times 10^{-08}$** | $2.8 \times 10^{-05}$ | $4.6 \times 10^{-01}$ | **$4.2 \times 10^{-12}$** | $7.9 \times 10^{-03}$ | **$4.2 \times 10^{-11}$** |
| | *PAH* | 12 | 6 | 27 | $4.3 \times 10^{-05}$ | $2.0 \times 10^{-01}$ | $5.3 \times 10^{-04}$ | **$7.2 \times 10^{-06}$** | $9.3 \times 10^{-03}$ | **$1.3 \times 10^{-06}$** |
| | *DHODH* | 16 | 5 | 90 | **$1.3 \times 10^{-07}$** | $5.3 \times 10^{-02}$ | $1.3 \times 10^{-02}$ | **$2.9 \times 10^{-06}$** | $7.6 \times 10^{-04}$ | **$7.7 \times 10^{-06}$** |
| | *BCAT2* | 19 | 3 | 133 | **$8.4 \times 10^{-06}$** | $4.0 \times 10^{-01}$ | $1.7 \times 10^{-02}$ | $1.1 \times 10^{-05}$ | $6.1 \times 10^{-03}$ | $3.7 \times 10^{-05}$ |
| | *ALDH1L1* | 3 | 8 | 103 | **$8.3 \times 10^{-08}$** | $1.7 \times 10^{-02}$ | $5.1 \times 10^{-03}$ | **$2.1 \times 10^{-06}$** | $1.7 \times 10^{-05}$ | $4.2 \times 10^{-05}$ |
| | *MED1* | 17 | 3 | 147 | $9.4 \times 10^{-01}$ | $1.8 \times 10^{-01}$ | $8.7 \times 10^{-05}$ | $2.1 \times 10^{-03}$ | $7.1 \times 10^{-01}$ | $3.4 \times 10^{-04}$ |
| | *STK33* | 11 | 6 | 180 | $4.8 \times 10^{-01}$ | $7.1 \times 10^{-01}$ | $7.0 \times 10^{-04}$ | $3.8 \times 10^{-02}$ | $6.4 \times 10^{-01}$ | $6.3 \times 10^{-03}$ |

To directly compare our results with existing methods we applied GAMuT, DKAT and MSKAT to the METSIM dataset. Since these methods cannot be applied to related individuals, we eliminated 1332 individuals that were related up to second degree, leaving us 7213 individuals. Table 4 shows p-values of different methods on the eight significant or suggestive genes displayed in Table 2. Since DKAT and GAMuT had nearly identical p-values when the same kernels were used, DKAT p-values were not shown in Table 4. For unrelated individuals, as expected, p-values produced by MSKAT with Q statistic, GAMuT with projection phenotype kernel and PhC (with SKAT Σ*_G_*) were very similar, and minP_com_ provided similar or more significant p-values than PhC. Interestingly MSKAT with Q’ statistic and GAMuT with linear phenotype kernel have less significant p-values than the other tests. We found that in 5 of the 8 genes in Table 4, using all individuals with kinship correction produced more significant PhC and minPcom p-values than using only unrelated individuals. Further, we have listed the top 10 genes for each of PhC, GAMuT and MSKAT with unrelated individuals (Supplementary Table S4). Except for the genes in Table 4, no other genes were found to be significant or suggestive.

**Table 4:** P-values for genes reported in Table 2 using MSKAT, GAMuT and Multi-SKAT (PhC and minP_com_). P-values in columns 2 to 7 were calculated with unrelated individuals (n = 7213), while those in columns 8 and 9 were calculated using all individuals (n = 8545). Both MSKAT (*Q* and *Q*′ statistic) and GAMuT (Projection and Linear phenotype kernel) p-values were calculated with the linear weighted genotype kernel. DKAT p-values were nearly identical to those of GAMuT when the same kernels were used (data not shown). P-values smaller than the Bonferroni corrected significance *α* = 9.6 × 10^−6^ were marked as bold.

| Gene | Unrelated individuals (n = 7213) |  |  |  |  |  | All individuals (n = 8545) |  |
| --- | --- | --- | --- | --- | --- | --- | --- | --- |
| | MSKAT<br>($Q$) | MSKAT<br>($Q'$) | GAMuT<br>(Projection) | GAMuT<br>(Linear) | PhC<br>($\Sigma_G = \text{SKAT}$) | minP<sub>com</sub> | PhC<br>($\Sigma_G = \text{SKAT}$) | minP<sub>com</sub> |
| *GLDC* | **$8.9 \times 10^{-54}$** | **$6.1 \times 10^{-15}$** | **0\*** | **$6.2 \times 10^{-15}$** | **$8.1 \times 10^{-54}$** | **$1.3 \times 10^{-53}$** | **$2.9 \times 10^{-73}$** | **$2.3 \times 10^{-72}$** |
| *HAL* | $9.5 \times 10^{-05}$ | $4.2 \times 10^{-02}$ | $9.6 \times 10^{-05}$ | $4.3 \times 10^{-02}$ | $9.5 \times 10^{-05}$ | **$2.9 \times 10^{-07}$** | **$2.9 \times 10^{-06}$** | **$9.5 \times 10^{-11}$** |
| *DHODH* | **$2.1 \times 10^{-06}$** | $2.2 \times 10^{-03}$ | **$2.4 \times 10^{-06}$** | $2.2 \times 10^{-03}$ | **$1.9 \times 10^{-06}$** | **$8.3 \times 10^{-06}$** | **$3.1 \times 10^{-08}$** | **$1.1 \times 10^{-07}$** |
| *PAH* | $9.9 \times 10^{-06}$ | $1.3 \times 10^{-02}$ | $1.0 \times 10^{-05}$ | $1.3 \times 10^{-02}$ | $9.9 \times 10^{-06}$ | $5.1 \times 10^{-05}$ | **$3.9 \times 10^{-07}$** | **$1.9 \times 10^{-06}$** |
| *MED1* | $1.7 \times 10^{-03}$ | $2.7 \times 10^{-01}$ | $1.7 \times 10^{-03}$ | $2.7 \times 10^{-01}$ | $1.7 \times 10^{-03}$ | $2.8 \times 10^{-05}$ | $2.8 \times 10^{-04}$ | **$4.9 \times 10^{-06}$** |
| *STK33* | $6.8 \times 10^{-04}$ | $6.9 \times 10^{-01}$ | $6.7 \times 10^{-04}$ | $6.9 \times 10^{-01}$ | $6.7 \times 10^{-04}$ | **$1.9 \times 10^{-06}$** | $1.1 \times 10^{-03}$ | $4.2 \times 10^{-05}$ |
| *ALDH1L1* | $5.9 \times 10^{-05}$ | $3.1 \times 10^{-02}$ | $6.1 \times 10^{-05}$ | $3.0 \times 10^{-02}$ | $6.0 \times 10^{-05}$ | **$9.5 \times 10^{-06}$** | $1.6 \times 10^{-04}$ | $5.4 \times 10^{-05}$ |
| *BCAT2* | $6.1 \times 10^{-04}$ | $1.5 \times 10^{-02}$ | $6.3 \times 10^{-04}$ | $1.5 \times 10^{-02}$ | $6.2 \times 10^{-04}$ | $3.7 \times 10^{-05}$ | $2.4 \times 10^{-03}$ | $6.2 \times 10^{-05}$ |

\* : GAMuT software collapses any p-value lower than $10^{-50}$ to 0

**Table 5:** Single phenotype SKAT-O with kinship adjustment test for the METSIM study data (n = 8545). P-values smaller than the Bonferroni corrected significance *α* = 9.6 ×10^−6^ were marked as bold.

| Gene | Ala | Gln | Gly | His | Ile | Leu | Phe | Tyr | Val |
| --- | --- | --- | --- | --- | --- | --- | --- | --- | --- |
| *GLDC* | $2.9 \times 10^{-01}$ | $2.9 \times 10^{-03}$ | **$2.5 \times 10^{-64}$** | $1.0 \times 10^{-02}$ | $1.5 \times 10^{-01}$ | $3.4 \times 10^{-02}$ | $8.5 \times 10^{-01}$ | $6.6 \times 10^{-01}$ | $4.7 \times 10^{-03}$ |
| *HAL* | $9.9 \times 10^{-01}$ | $1.1 \times 10^{-01}$ | $3.1 \times 10^{-01}$ | **$3.2 \times 10^{-09}$** | $2.6 \times 10^{-01}$ | $2.5 \times 10^{-01}$ | $6.6 \times 10^{-01}$ | $1.2 \times 10^{-01}$ | $3.5 \times 10^{-01}$ |
| *DHODH* | **$1.4 \times 10^{-07}$** | $9.9 \times 10^{-01}$ | $9.0 \times 10^{-02}$ | $3.6 \times 10^{-01}$ | $7.7 \times 10^{-01}$ | $1.7 \times 10^{-01}$ | $1.9 \times 10^{-01}$ | $3.0 \times 10^{-01}$ | $1.0 \times 10^{-02}$ |
| *PAH* | $7.9 \times 10^{-01}$ | $3.0 \times 10^{-01}$ | $2.8 \times 10^{-01}$ | $8.6 \times 10^{-01}$ | $4.5 \times 10^{-01}$ | $8.1 \times 10^{-01}$ | $6.8 \times 10^{-05}$ | $4.0 \times 10^{-01}$ | $9.9 \times 10^{-01}$ |
| *MED1* | $1.0 \times 10^{-01}$ | $5.1 \times 10^{-01}$ | $7.9 \times 10^{-01}$ | $5.9 \times 10^{-01}$ | $3.3 \times 10^{-01}$ | $1.4 \times 10^{-01}$ | $6.9 \times 10^{-01}$ | $6.8 \times 10^{-02}$ | $6.7 \times 10^{-01}$ |
| *STK33* | $8.4 \times 10^{-01}$ | $8.2 \times 10^{-01}$ | $3.5 \times 10^{-01}$ | $5.7 \times 10^{-01}$ | $6.6 \times 10^{-01}$ | $1.0 \times 10^{-01}$ | $9.9 \times 10^{-01}$ | $8.1 \times 10^{-01}$ | $4.1 \times 10^{-01}$ |
| *ALDH1L1* | $6.7 \times 10^{-01}$ | $7.6 \times 10^{-01}$ | **$9.3 \times 10^{-09}$** | $7.3 \times 10^{-01}$ | $3.4 \times 10^{-01}$ | $3.2 \times 10^{-01}$ | $9.9 \times 10^{-01}$ | $5.9 \times 10^{-01}$ | $2.5 \times 10^{-01}$ |
| *BCAT2* | $1.4 \times 10^{-01}$ | $2.6 \times 10^{-01}$ | $8.2 \times 10^{-01}$ | $9.9 \times 10^{-01}$ | $1.0 \times 10^{-01}$ | $2.1 \times 10^{-02}$ | $5.4 \times 10^{-01}$ | $5.0 \times 10^{-01}$ | **$1.2 \times 10^{-06}$** |

Overall, our METSIM amino acid data analysis suggests that the proposed method can be more powerful than the single phenotype tests as well as existing tests, while maintaining type I error rate even in the presence of the relatedness. It also shows that the omnibus tests (minP and minPcom) provides robust performance by effectively aggregating results of various kernels.

### Computation Time

When Σ*_P_* and Σ*_G_* are given, p-values of Multi-SKAT are computed by the Davies method (Davies, 1980), which inverts the characteristic function of the mixture of chi-squares. On average, Multi-SKAT tests for a given Σ*_P_* and Σ*_G_* required less than 1 CPU sec (Intel Xeon 2.80 GHz) when applied to a dataset with 5000 independent individuals, 20 variants and 10 phenotypes (Supplementary Table S1). With the kinship adjustment for 5000 related individuals, computation time was increased to 3 CPU sec. Since minP_com_ requires only a small number of resampling steps to estimate the correlation among tests, it is still scalable for genome-wide analysis. In the same dataset, minP_com_ required 4 and 10 CPU sec on average without and with the kinship adjustment, respectively. Further, Multi-SKAT given Σ*_P_* and Σ*_G_*, is computationally equivalent to MSKAT and takes less than 1 CPU-sec for up to 20,000 samples, with 20 variants (Supplementary Figure S7 A), while GAMuT takes considerably more time than these two. The performance of minP_com_ is similar to GAMuT for small and moderate sample sizes (7.5 and 7.1 CPU-secs respectively for 10,000 samples) and performs better than GAMuT for larger sample sizes (14.9 and 34.6 CPU-secs respectively for 20,000 samples). Computation time of all the methods were slightly increased when the number of variants were 100 (Supplementary Figure S7 B). Analyzing the METSIM dataset with minP_com_ required 10 hours when parallelized into 5 processes.

## Discussion

In this article, we have introduced a general framework for rare variant tests for multiple phenotypes. As demonstrated, Multi-SKAT gains flexibility with regard to modeling the relationship between phenotypes and genotypes through the use of the kernels Σ*_P_* and Σ*_G_*. Many published methods, including GAMuT, MSKAT and MAAUSS, can be viewed as special cases of the Multi-SKAT test with corresponding values of Σ*_P_* and Σ*_G_*, which illuminates the underlying assumptions of these methods and their relationships. In addition, by unifying existing methods to the common framework, our approach provides a way to combine different methods through the minimum p-value based omnibus test. Our method can also adjust for sample relatedness. From simulation studies we have found that the proposed method is scalable to genome-wide analysis and can outperform the single phenotype test and existing multiple phenotype tests. The METSIM data analysis demonstrated that the proposed methods perform well in practice.

It is natural to assume that different genes follow different models of association. For some genes, the effect of the variants on the phenotypes might be independent of each other, thus best detected by the Het phenotype kernel for Σ*_P_*, while for others, the effects might be nearly the same and best detected by the Hom phenotype kernel. If the kernel structures are chosen based on prior knowledge and the selected Σ*_G_* and Σ*_P_* do not reflect underlying biology, the test may have substantially reduced power. The omnibus test, which uses the minimum p-value from the various choices of kernels, has been a useful approach under such situations in genetic association analysis (Lee et al., 2012b; Urrutia et al., 2015; Zhan et al., 2017). We applied this ominibus test to Multi-SKAT and used a Copula to obtain p-values. As seen in simulation studies and real data analysis, our omnibus approaches (minP and minPcom) are scalable to genome-wide analysis and provide robust power regardless of underlying genetic models.

Multi-SKAT retains most of the desirable properties of SKAT. The asymptotic p-values of all the MultiSKAT tests, other than minP and minPcom, can be analytically obtained via Davies’ method. The p-value calculations for minP and minP_com_ depend on a resampling based approach but a reliable estimate can be obtained using a small number of resampling steps. Thus, computationally all the Multi-SKAT tests are scalable at the genome-wide level. This method also allows the inclusion of prior information through weighting of variants.

Additionally, Multi-SKAT can adjust for the relatedness among study individuals by accounting for their kinship matrix. As shown in Supplementary Figure S5, in the presence of related individuals, lack of adjustment for relatedness can produce inflated type I error rate. Since Multi-SKAT is a regression based approach, it effectively incorporates the relatedness by including a random effect term for kinship. Type I error simulation and METSIM data analysis show that our approach produced more significant p-values than alternative methods, like GAMuT and MSKAT, while controlling type I error rates.

Although Multi-SKAT provides a general framework for gene-based multiple phenotype tests, the current approach is limited to continuous phenotypes. In the future, using a generalized mixed effect model framework, we aim to extend Multi-SKAT to binary phenotypes.

In summary, we have developed a powerful multiple phenotype test for rare variants. The proposed method has robust power regardless of the underlying biology and can adjust for family relatedness. Our method can be a scalable and practical solution to test for multiple phenotypes and will contribute to detecting rare variants with pleiotropic effects. All our methods are implemented in the R package MultiSKAT (see Web Resource).

## Web Resources

MultiSKAT R-package: https://github.com/diptavo/MultiSKAT

GAMuT R-package: https://epstein-software.github.io/GAMuT

MSKAT R-package: https://github.com/baolinwu/MSKAT

PHENIX R-package: https://mathgen.stats.ox.ac.uk/genetics_software/phenix/phenix.html

Online Mendelian Inheritance in Man (OMIM): http://www.omim.org

## Acknowledgments

This work was supported by grants R01 HG008773 and R01 LM012535 (D.D. and S.L.) and R01 HG000376 and U01 DK062370 (L.S. and M.B.) from the National Institutes of Health. We would like to thank investigators of the METSIM study for access to the genotype and amino-acid phenotype data.

## Appendices

## A. Principal Component (PC) Kernel

Let *L_i_* be the loading vector for the *i^th^* PC, which produces the *i^th^* PC score *P_i_* = *YL_i_*. In PCA-based analysis, PC scores are used as outcomes instead of original *Y*. Since the genetic information regarding the phenotypes may not be confined to the top few PCs (Aschard et al., 2014), we first consider using all PCs. Let *P* = (*P_i_*, ⋯ *P_K_*). Since PCs are orthogonal, we assume genetic effects to multiple PCs are heterogeneous, which resulted in

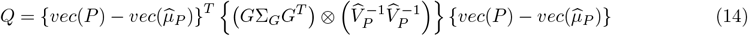

where 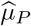 is the mean of *P* under the null hypothesis and 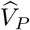 is the estimated covariance matrix between the PC’s. 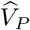 will be a diagonal matrix since PCs are orthogonal. Equation (14) can be written as

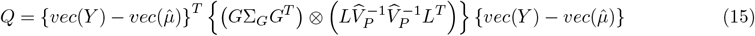

where *L* = (*L*_1_, ⋯, *L_K_*) is a *K × K* PC loading matrix. Equation (15) shows that by using 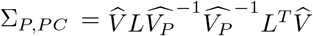, we can carry out PC-based tests. It is to be noted that the genetic effects of the PC’s do not need to be assumed to be heterogeneous. Any kernel structure that is applicable to the test statistic in equation 4 can be applied here as well.

## B. Relationship between Multi-SKAT and existing methods

For the ease of algebraic expressions, we will consider that all the *K* phenotypes have residual variance 1. For the general case of different residual variances, Σ*_P_* should be replaced by 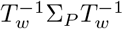 where *T_w_* = *diag*(*σ*_1_, ⋯, *σ_K_*), *σ_k_* being the residual standard error of *k^th^* phenotype.

### B.1. MSKAT

The *Q* statistic of MSKAT(Wu and Pankow, 2016) is given by

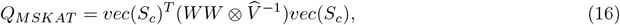

where 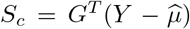 is a matrix of score statistics (Wu and Pankow, 2016). Using row-vectorization properties

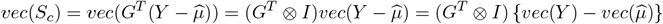

Then *Q_MSKAT_* can be written as

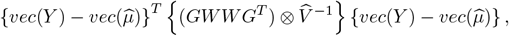

which is the Multi-SKAT test statistics with Σ*_G_* = *WW* and 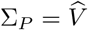. Further, the *Q*′ of MSKAT is given by

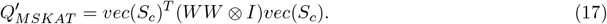

Using the similar algebra as above, this can be written as

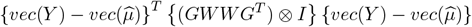

which is the Multi-SKAT test statistics with Σ*_G_* = *WW* and 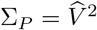.

### B.2. GAMuT

Suppose 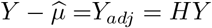 and *G_adj_* = *HG* are covariate adjusted phenotype and genotype matrices where *H* = *I − X*(*X^T^X*)^−1^*X^T^*. With the intercept in *X*, *Y_adj_* and *G_adj_* are mean centered. The covariate adjusted GAMuT test statistics is

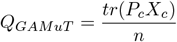

where

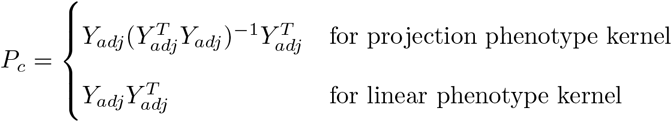

and 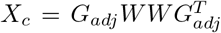. Using the fact that 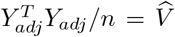 is the estimate of variance after adjusting covariates and 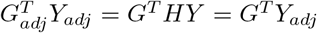 (since *H* is a symmetric idempotent matrix), we show, for the projection kernel

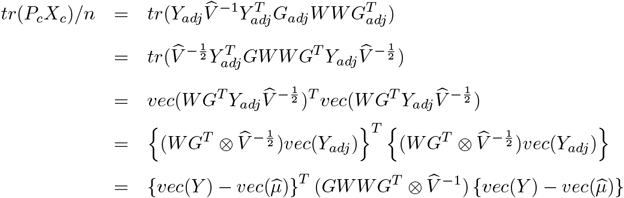

which is the same as the Multi-SKAT test statistic with Σ*_G_* = *WW* and 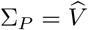. Similarly for the linear kernel,

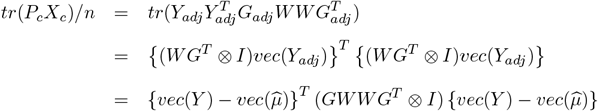

which is the Multi-SKAT test statistic with Σ*_G_* = *WW* and 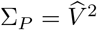.

### B.3. MAAUSS and MF-KM

There exists two different version of the MAAUSS tests. The homogeneous version of MAAUSS assumes that the effects of a variant on multiple phenotypes are identical and uses the following test statistic

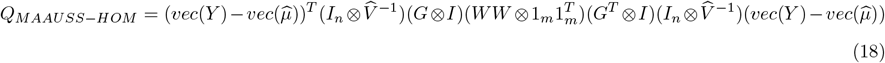

which is identical to the Multi-SKAT test statistic with Σ*_G_* = *WW* and 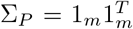. The heterogeneous version of MAAUSS assumes that the effects of a variant on multiple phenotypes are independent, and uses the following test statistic

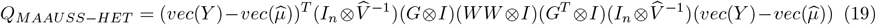

which is identical to the Multi-SKAT test statistic with Σ*_G_* = *WW* and 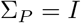. Note that the test statistic of MF-KM is exactly the same as *Q_MAAUSS-HET_*.

## C. Backward elimination procedure to identify associated phenotypes

After identifying the gene or region associated with multiple phenotypes, next question would be identifying truly associated phenotypes. Here we present a simple backward elimination algorithm to iteratively remove relatively less important phenotypes. A similar method has previously been applied to identify rare causal variants in an associated gene (Ionita-Laza et al., 2014).

- Step 1. Start with a set of *k* phenotypes *Phen_Current_* = {*y*_1_, *y*_2_, ⋯ *y_k_*} and compute a Multi-SKAT test association p-value for the set *Phen_Current_* denoted by *p_Current_*.
- Step 2. Remove each of the phenotypes one at a time from the set *Phen_Current_*. The resulting set is 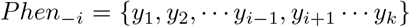 for *i* = 1, 2, ⋯, *k* and compute the corresponding p-values *p_−i_* for that same Multi-SKAT test.
- Step 3. Remove the phenotype *j* that leads to the smallest p-value, i.e. *j* = *argmin*{*p*_−1_, *p*_−2_, ⋯, *p_−k_*}. Update *Phen_Current_* to *Phen_−j_*.
- Step 4. Continue removing phenotypes till only 1 phenotype is left.

Supplementary Table S2 shows the backward elimination results of 5 most significant and suggestive genes in the METSIM study data analysis as per the p-values reported by minP_com_. Although this procedure does not provide a set of phenotypes truly associated, it provides the relative importance of the phenotypes in driving association signals. For example, the minP_com_ p-value for *GLDC* was 2.3 × 10^−72^. When each of the phenotypes were removed one at a time and the minPcom p-values were calculated on the remaining 8 phenotypes, we found that eliminating Isoleucine (Ile) actually improved the signal. The minP_com_ p-value of the set of 8 amino acids leaving out Ile was 2.8 × 10^−73^. This indicates that Isoleucine has very minimal contribution to the association between the amino acids and *GLDC*. Subsequently, Valine was the next phenotype to be eliminated indicating that it has the next lowest contribution after Isoleucine. Carrying out this procedure further, we find that Glycine is the last phenotype to remain indicating that it is the strongest driver of the signal. This is in agreement to the single phenotype SKAT-O results (Table 5). Similary for genes *HAL*, *DHODH*, *PAH* and *MED1*, Histine, Alanine, Phenylalanine and Tyrosine were the most associated phenotypes, respectively. Interestingly for *PAH* and *MED1*, single phenotype p-values are not significant, which suggests that multiple phenotypes are associated with these genes.

